# Cognitive and motor perseveration are associated in older adults

**DOI:** 10.1101/2020.09.17.302083

**Authors:** Carly Sombric, Gelsy Torres-Oviedo

## Abstract

Aging causes perseveration (difficulty to switch between actions) in motor and cognitive tasks, suggesting that the same neural processes could govern these abilities in older adults. To test this, we evaluated the relation between independently measured motor and cognitive perseveration in young (21.4±3.7 y/o) and older participants (76.5±2.9 y/o). Motor perseveration was measured with a locomotor task in which participants had to transition between distinct walking patterns. Cognitive perseveration was measured with a card matching task in which participants had to switch between distinct matching rules. We found that perseveration in the cognitive and motor domains were positively related in older, but not younger individuals, such that participants exhibiting greater perseveration in the motor task also perseverated more in the cognitive task. Additionally, exposure reduces motor perseveration: older adults who had practiced the motor task could transition between walking patterns as proficiently as naïve, young individuals. Our results suggest an overlap in neural processes governing cognitive and motor perseveration with aging and that exposure can counteract the age-related motor perseveration.

**Highlights:**

- Movement carryover from the treadmill to overground indicates motor perseveration.
- Greater motor and cognitive perseveration are associated in older adults.
- Motor perseveration in older adults can be reduced with practice.
- New motor memories are similarly forgotten in older and younger adults.

## 1.1 Introduction

It is important for the motor system to develop context-specific motor memories that benefit future performance. For example, humans improve their ability to walk on an icy terrain by practicing two elements of this gait pattern : 1) by perfecting a walking pattern to increase stability on the slippery surface and 2) by improving their ability to switch between context-specific walking patterns once the terrain changes. In this example of walking on ice, it is easy to imagine that failure to switch walking patterns, a form of motor perseveration, could lead to falls or inefficient gait. Notably, failure to accommodate new toe clearance (Bunterngchit, Yuthachai, Thurmon Lockhart, Jeffrey C. Woldstad, 2000) or friction demands of the walking surface (Lockhart et al., 2002; Lockhart, 1997) contributes to fall risk in the elderly. Despite the drastic impact on older population’s daily living, little is known about neural processes governing the age-related decline of motor switching (i.e., increased motor perseveration). It is also observed that healthy aging induces greater perseveration in cognitive tasks requiring participants to switch strategies (Albergaria et al., 2018; Boone et al., 1993; Daigneault et al., 1992; Haaland et al., 1987; Head et al., 2009; Kray and Lindenberger, 2000; Ridderinkhof et al., 2002; Volkow et al., 1998) or decisions (Eppinger et al., 2011). Pathways in the basal ganglia regulate motor switching in the young nervous system (Balser et al., 2014; Brown and Almeida, 2011; Leunissen et al., 2013). However, prefrontal neural resources, usually involved in cognitive tasks, are recruited for motor switching with healthy aging (Coxon et al., 2010) presumably to compensate for deteriorated basal ganglia function in the elderly (Bäckman et al., 2006; Ota et al., 2006; Walhovd et al., 2011). Thus, neural processes leading to perseveration in motor and cognitive domains might become more unified as we age.

In this study, we tested the possibility that the same underlying mechanisms mediate age-related changes in motor and cognitive perseveration. This is feasible given extensive evidence of a direct associations between motor and cognitive processes in older adults (Anguera et al., 2011; Bo et al., 2009; Langan and Seidler, 2011; Trewartha et al., 2014; Wolpe et al., 2020). Consistently, it has been shown that interventions that reduce motor perseveration also diminish cognitive perseveration in the elderly (Coubard et al., 2011), which suggests that processes underlying motor and cognitive switching are indeed linked in older populations. Interestingly, cognitive switching appears to interfere, rather than favor, motor switching in the context of locomotion (Sombric et al., 2017). More specifically, we found that older individuals proficient at switching actions in response to explicit instructions in a cognitive task, had difficulties at switching between motor patterns that are controlled implicitly, such as timing between steps (Sombric et al., 2017). This finding suggests that cognitive-mediated processes for switching are recruited to transition between motor patterns in older individuals, but that this is only beneficial when said patterns are controlled explicitly.

We tested the hypothesis that cognitive switching processes influence motor switching in older adults. However, this relation is only beneficial in locomotion when switching between spatial motor patterns (i.e., “where” to step), which are more explicitly controlled than temporal motor patterns (i.e., “when” to step). To test this hypothesis, young and older adults adapted their walking pattern on a split-belt treadmill that drives the legs at different speeds. We subsequently measured motor perseveration as either the participant’s difficulty to disengage spatial (i.e., explicitly controlled) or temporal (i.e., implicitly controlled) aspects of the split-belt gait pattern when transitioning to walking overground. We also measured cognitive perseveration with a card matching task in which participants had to switch between different matching rules. We anticipated that cognitive perseveration would be associated to motor perseveration of the spatial motor pattern, but not the temporal one. These findings would support the idea that cognitive processes are recruited to compensate for age-related decline in motor switching, but this compensation would only benefit motor aspects controlled explicitly. In a post hoc analysis we evaluated the effect of repeated exposure to the split-belt task onto motor perseveration in older adults. This was done given the unexpected observation that older individuals, who had previously experienced the split-belt task, exhibited less motor perseveration than naïve, younger participants. Taken together, our findings indicate the extent to which cognitive-mediated switching and practice can help older adults regain motor switching abilities similar to younger individuals.

## 1.2 Methods

### 1.2.1 Subjects

We assessed motor and cognitive perseveration in a group of young (n=11, #women=7, 21.4±3.7 y/o) and older adults (n=11, #women=6, 76.5±2.9 y/o). Motor perseveration was evaluated with a locomotor task in which participants had to transition between distinct walking environments (i.e., split-belt treadmill with legs moving at different speeds vs. overground walking). Cognitive perseveration was evaluated with a card matching task in which participants had to switch between matching rules (i.e., modified Wisconsin Card Sorting Test). The Institutional Review Board at the University of Pittsburgh approved the experimental protocol and all subjects gave informed consent prior to testing.

### 1.2.2 General Paradigm

The general protocol consisted of paradigms in the motor and cognitive domains. The motor paradigm enabled us to quantify age-related differences in acquisition of a split-belt motor pattern and switching between said pattern and regular overground walking. The cognitive paradigm consisted of two tests: one aimed to quantify age-related differences in cognitive perseveration and a second one to measure spatial working memory, which is a cognitive ability known to be strongly related to motor learning processes (Anguera et al., 2011; Bo et al., 2009; Langan and Seidler, 2011; Trewartha et al., 2014; Uresti-Cabrera et al., 2015). We evaluated the association between each of these cognitive abilities and motor perseveration to determine if motor switching was specifically associated to cognitive switching or to better cognition in general.

#### 1.2.2.1 Locomotor Paradigm

All subjects completed the same locomotor paradigm consisting of 5 epochs: Baseline, Adaptation, Catch, Re-Adaptation, and Post-Adaptation (Figure 1A). The Baseline epoch was used to characterize each subject’s baseline gait overground and on the treadmill. In the overground condition, subjects walked back and forth on a walkway (approximately 7m-long) for 4 minutes (approx. 100 strides) at a self-selected pace. In the treadmill condition, subjects walked for 150 strides at 0.75 m/s. The Adaptation epoch consisted of 600-strides of split-belt walking in which the (self-reported) dominant leg walked at 1.00 m/s (i.e., fast leg) and the other leg walked at 0.50 m/s (i.e., slow leg). Subjects walked in this condition for 600 strides to ensure that a steady state behavior was reached in all individuals. All subjects, young and old, took sitting breaks every 150 strides. Each break in the young group lasted about 8 minutes and 10 seconds, which was the average duration of the sitting breaks in the old group. We imposed similar break durations between the age groups to determine if age-related differences in forgetting (i.e., decay of the split-belt pattern due to the passage of time) reported in previous studies (Malone and Bastian, 2014; Sombric et al., 2017) were due to group differences in break durations. The Catch epoch consisted of 10 strides of tied walking at 0.75 m/s to measure After-Effects on the treadmill (i.e., context in which subjects acquired the split-belt pattern). The Re-Adaptation epoch consisted of 300 strides of split-belt walking at the same speed as the Adaptation epoch. This was done such that all individuals again reached a steady state pattern in the split-belt condition before walking overground. The Post-Adaptation epoch consisted of 6 minutes (approximately 150 strides) of walking overground to evaluate each subject’s ability to disengage the split-belt walking pattern when transitioning into a different environment. Subjects started on the treadmill and walked back and forth at a self-selected speed on the same walkway as in the Baseline epoch. A stride is defined as the duration from one heel-strike to the subsequent heel-strike with the same leg. Strides were counted on the treadmill in real-time to ensure that the Baseline, Adaptation, and Re-Adaptation epochs contained the same number of strides across individuals walking with different cadences.

**Figure 1:**
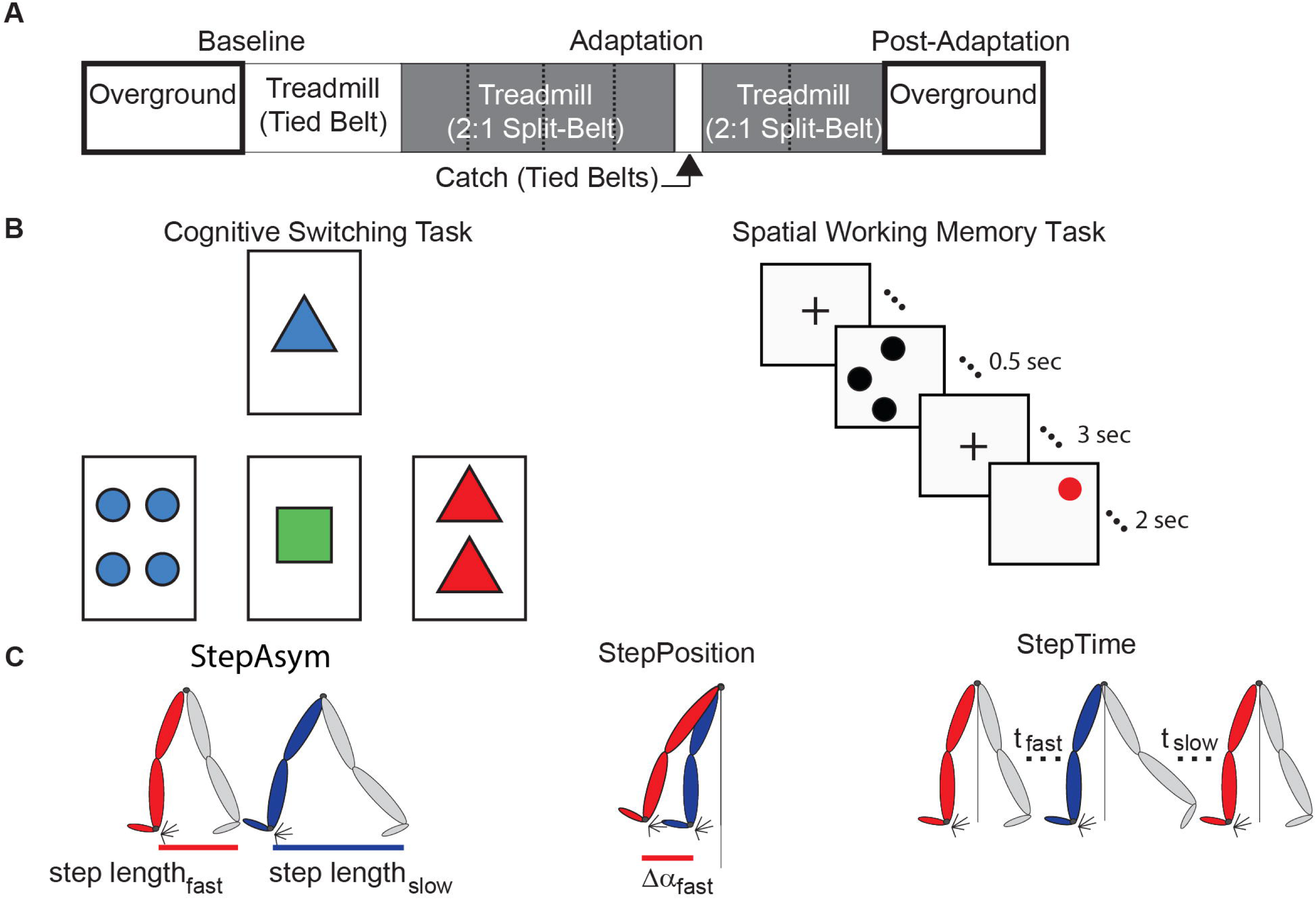
Experimental paradigms and parameter definitions. **(A)** The split-belt treadmill paradigm used for both age groups is illustrated. Resting breaks, when subjects did not walk, are indicated by dashed lines. These were taken every 150 strides and lasted 8 minutes and 10 seconds on average. **(B)** The left panel is a sample screen for the card matching test to assess cognitive perseveration. The right panel demonstrates the temporal progression of a single trial of the computer-based test to assess spatial working memory accuracy. **(C)** This schematic adapted from Finley et al., (2015) illustrates Step Length Asymmetry (StepAsym) which can be decomposed into gait aspects that are controlled more explicitly (StepPosition) or implicitly (StepTime).

#### 1.2.2.2 Cognitive Paradigm

Older adults first underwent cognitive screening to rule out cognitive disability or decline. Specifically, intelligence quotients (IQ) were estimated with The Wechsler Test of Adult Reading (WTAR) to estimate premorbid cognitive ability and The Modified Mini-Mental State (3MS) examination was used to screen for dementia. All participants scored within normal limits for both IQ and general cognitive ability (WTAR range = 89-125; 3MS range = 86-99 (/100)) (Li et al., 2016).

Then, two cognitive abilities were assessed in all subjects: cognitive switching and spatial working memory. The assessment of these cognitive abilities allowed us to determine the specificity of a possible association between motor switching and distinct cognitive processes.

Cognitive perseveration was evaluated with a computer-based task inspired by the Wisconsin Card Sorting Test. In brief, participants had to match cards by either count, color, or shape. These three matching rules were given to the participants before the task started. More specifically, 4 electronic cards were displayed with a specific count (1, 2, 3, or 4 items) of colored (red, blue, green, or yellow) shapes (squares, circles, triangles, or plus signs) on a computer monitor (Figure 1B, left panel). For each trial, a single reference card was displayed at the top of the screen and three test cards were displayed below. Each of the test cards matched the reference card in only one of the three possible matching rules (Figure 1B, left panel; count, shape, or color). Unlike a traditional Wisconsin Cards Sorting Test, the experimenter demonstrated the matching test and participants had a familiarization period during which they practiced matching cards using each of the matching rules and switching between them. This familiarization was done as in previous work (Stuss et al., 2000) to obtain a more robust assessment of age-related differences in people’s ability to switch choices (Nelson, 1976). More specifically, the experimenter first explained the matching game. Then, a baseline matching period was collected for each matching rule to measure subjects’ baseline accuracy when the matching rule was not changing. This baseline period was collected for each matching rule and it consisted of two untimed practice trials followed by 64 timed trials (3 seconds to respond) with a single matching rule. The word “correct” or “incorrect” appeared for 0.5 seconds after each trial indicating if the match was correct or incorrect. Next, we tested subjects’ ability to switch between matching rules based on feedback on their matching accuracy. Subjects were told that there were three possible matching rules and that the matching rule would change unexpectedly throughout the experiment. Specifically, “Sometimes you will be matching by count, sometimes by shape, and sometimes by color. The computer will not tell you which matching rule you should match with. You will have to determine the rule by trial-and-error. If you get a match correct, you should continue to use this matching rule. If you get a match incorrect you should try a different matching rule”. Subjects performed untimed practice trials during which they matched cards under the supervision of the experimenter to familiarize themselves with the task. This ensured that both older and younger adults understood the task; so that any differences in behavior would be due to cognitive constraints, rather than poor understanding of the computer-based test. Following these practice trials, subjects performed the actual test which consisted of a total of 128 matching trials. Subjects had 5 seconds to respond to each match and feedback on their match (i.e., correct or incorrect) was displayed for 1 second. The rule changed in a predefined order after 3-5 consecutive correct matches.

We also assessed subjects’ spatial working memory because this is a measure of cognition that is broadly related to physical fitness (Erickson et al., 2011, 2009), integrity of neural networks associated with motor learning (Hötting et al., 2013; Salmi et al., 2018), and behavioral motor learning outcomes (Anguera et al., 2011; Bo et al., 2009; Christou et al., 2016; Langan and Seidler, 2011; Trewartha et al., 2014; Uresti-Cabrera et al., 2015; Vandevoorde and Orban de Xivry, 2019). Spatial working memory was assessed with a spatial working memory task similar to previous work (e.g., Erickson et al., 2012, 2011, 2009; Weinstein et al., 2012) (Figure 1B, right panel). Each of the 45 trials started with a fixation mark in the middle of the screen for 1 second. Next, 1-3 black dots appear on the screen for 0.5 seconds, followed by a 3 second hold phase where the fixation mark again appeared. Finally, a single red dot appeared for 2 seconds and subjects indicated if the red dot was in the same position as one of the previously seen black dots. The results from this task were not recorded in two out of the eleven younger participants due to technical difficulties. Thus, all results from the spatial working memory task consist of the measures on only nine, rather than eleven, young subjects.

### 1.2.3 Data Collection

#### 1.2.3.1 Locomotor Task

Kinematic data were collected to characterize subjects’ locomotor movements on the treadmill and overground. A motion analysis system (Vicon Motion Systems, Oxford, UK) was used to collect kinematic data at 100 Hz. A quintic spline interpolation was used to fill gaps in the raw kinematic data (Woltring; Vicon Nexus Software, Oxford, UK). Subjects’ movements were tracked via passive reflective markers placed bilaterally over the hip (greater trochanter) and ankle (lateral malleoulous) and asymmetrically on the thigh and shank to distinguish the legs. The duration of treadmill trials was defined by real time kinetic detection of heel strikes whereas the duration of overground trials was defined by elapsed time. Heel strikes were identified with raw vertical kinetic data collected from the instrumented treadmill (Bertec, Columbus, OH, United States). Given that force data was only available during treadmill trials, but not overground trials, the greatest forward excursion of the ankle was used to identify heel strikes post-processing so that the same heel strike detection could be used across treadmill and overground walking epochs as had been previously done (e.g., Sombric et al., 2017; Torres-Oviedo and Bastian, 2012, 2010).

#### 1.2.3.2 Cognitive Tasks

We used pen and paper versions of the WTAR and 3MS tests, whereas the other cognitive tests were administered with custom code created using the E-Prime Software Suite (Psychology Software Tools, Sharpsburgh, PA).

### 1.2.4 Data Analysis

#### 1.2.4.1 Locomotor Parameters

Spatial and temporal features of gait were characterized to determine if cognitive processes were distinctly associated with more explicitly controlled spatial motor patterns (i.e., “where” to step) or with more implicitly controlled temporal motor patterns (i.e., “when” to step) (Malone and Bastian, 2010). These locomotor parameters quantify the asymmetries between fast and slow legs’ movements on consecutive steps. Step Length Asymmetry (StepAsym) is a robust and clinically relevant measure (e.g., Reisman et al., 2013) conventionally used to characterize gait changes in split-belt adaptation studies (e.g., Reisman et al., 2005). StepAsym is defined as the difference in consecutive step lengths where step lengths are defined as the distance between the ankles at forward leg heel strike (Equation 1). Therefore, zero values indicate that both steps are of the same length, and positive values indicate that the step length of the leg that walks fast during Adaptation is longer and vice versa for negative values.

StepAsym was further decomposed into spatial and temporal asymmetries because subjects exhibit distinct motor perseveration in these two domains (Mariscal et al., 2020; Sombric et al., 2017)(Figure 1C). The decomposition of StepAsym in spatial and temporal parameters is described in detail by Finley and colleagues (Finley et al., 2015). In brief, spatial asymmetry, labeled as StepPosition, was characterized as the difference in the forward position of the legs (Equation 2), where Δα represents the difference in the forward position of the ankles at heel strike relative to the body (averaged position of the hip markers). Temporal asymmetry, labeled as StepTime, was characterized as the difference in the durations of each step (Equation 3), where t represents the time between one heel strike and the following heel strike of the other leg. To be consistent with prior work, StepTime was multiplied by the average step speed to convert to units of distance. StepVelocity (v) is a proxy for the belt speeds computed as the speed of the stance ankle relative to the hips (Equation 4). Finally, all parameters were normalized by Stride Length (SL), or the distance traveled by the ankle between consecutive heel strikes of the same ankle, so that all measures were unitless and therefore comparable across subjects taking different step sizes. These parameters are smoothed with a 5-stride running average for visualization purposes.

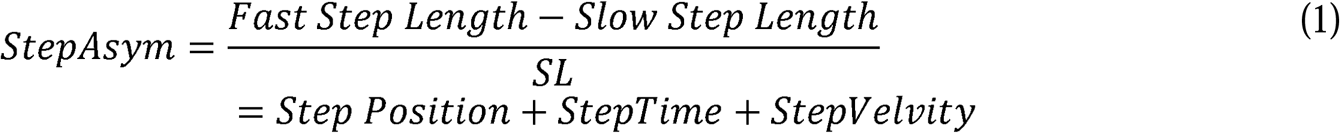

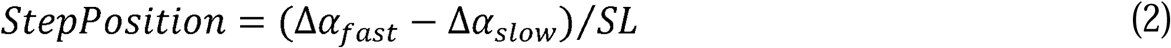

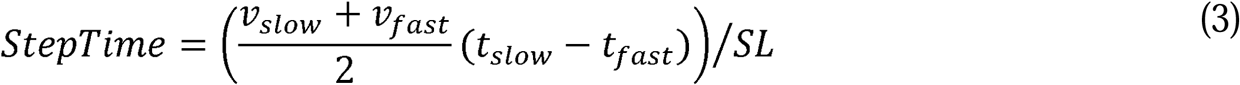

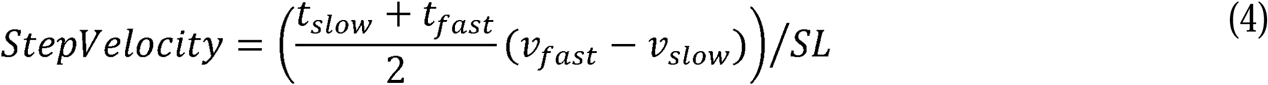

### 1.2.5 Outcome Measures

#### 1.2.5.1 Locomotor Outcome Measures

We used the same outcome measures as in our previous work (Sombric et al., 2017) to characterize the acquisition of the split-belt pattern and switching between said split-belt pattern and regular overground gait. These outcome measures were computed for each of the locomotor parameters described above (StepAsym, StepPosition, and StepTime).

The acquisition of the split-belt pattern was characterized with four measures: Steady State, Rate of Adaptation, %Forgetting, and Treadmill After-Effects. Steady State quantified how much subjects adapted each gait parameter in response to the split-belt perturbation. This was measured immediately before switching to walking overground during the Post-Adaptation epoch. Steady State was calculated as the difference between the average of the last 40 strides in the Re-Adaptation epoch and the baseline gait on the treadmill prior to the split-belt perturbation. The Rate of Adaptation was characterized with a time constant (τ). τ indicated the stride number at which subjects had achieved 63.2% of their Steady State value for the data smoothed with a 20-stride running average. Thus, a larger τ indicates slower adaptation than a small τ. In addition, %Forgetting characterized the decay of the split-belt pattern due to the passage of time during resting breaks. Large values of %Forgetting indicated that the split-belt pattern was very susceptible to the passage of time, whereas small values of %Forgetting indicated that this motor memory persisted over the duration of the resting break. %Forgetting was computed as the average change in the value for every gait parameter after (I_i_), relative to before (F_i_), each of the 3 resting breaks experienced during the Adaptation epoch (Equation 5). This difference was expressed as a percentage of the motor value before the break (F_i_).

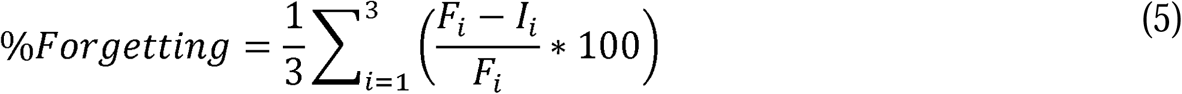

Lastly, after-effects on the treadmill were measured to quantify the extent to which participants maintained the split-belt pattern after the split perturbation was removed. These after-effects were measured in the same context in which the split-belt pattern was acquired (i.e., the treadmill). Consistently, After-Effects were quantified during the Catch epoch, when the split condition was briefly removed during the Adaptation period. More specifically, After-Effects were defined as the change in gait parameters during the first three strides of the Catch epoch relative to the Baseline epoch.

Motor perseveration was quantified as the after-effects observed when participants transitioned to walking overground following the Adaptation epoch. Notably, large after-effects overground indicated that subjects could not disengaged the split-belt pattern when walking on a different environment (i.e., poor motor switching performance). These Motor Perseveration Errors were computed as the average of the initial 5 steps Post-Adaptation during overground relative to Baseline overground. We also computed %Motor Perseveration, which is defined as Motor Perseveration Errors expressed as a percentage of the Steady State value reached by every individual. This was done to account for any potential differences in the split-belt pattern across participants. Note that this measure was not computed for StepAsym since Steady State of StepAsym approach zero, and therefore %Motor Perseveration for this parameter is numerically unstable.

#### 1.2.5.2 Cognitive Outcome Measures

Two outcome measures were computed to characterize subjects’ cognitive perseveration and spatial working memory. Cognitive Perseveration Errors were computed as the total number of matches that were made based on a previous matching rule, as done in published reports (Head et al., 2009). Thus, a large value for Cognitive Perseveration Errors indicated that individuals were poor at switching their actions in the cognitive switching task. On the other hand, spatial working memory ability was quantified with a measure of accuracy called Spatial Working Memory Accuracy. More specifically, accuracy was computed as the number of correct responses over the total number of responses in the spatial working memory task, which is consistent with prior work (Erickson et al., 2011; McAuley et al., 2011; Szabo et al., 2011; Voss et al., 2010).

### 1.2.6 Statistical Analysis

#### 1.2.6.1 Planned Analysis

Non-parametric statistics were utilized given the heterogenous nature of the older adults and the persistent violation of normality according to the Kolmogorov-Smirnov test. A Wilcoxon rank sum test was used to identify differences between the outcome measures (e.g., Motor Perseveration Errors, %Motor Perseveration, etc.) of younger and older adults. Cohen’s d effect sizes (e.s.) were computed for each of these statistics. Also, Spearman’s rank correlations were used to determine the extent to which motor switching was related to each of the cognitive outcome measures (i.e., Cognitive Perseveration Errors and Spatial Working Memory Accuracy). This was done to test our hypothesis that motor and cognitive switching are positively related in older adults; and that this relation between motor and cognitive domains is specific to switching and not cognitive performance in general. We performed linear correlations when Spearman’s rank correlations had significant rho’s. This was done for display purposes and consistency with prior literature (Sombric et al., 2017). We used a significance level of α = 0.05 for all the analyses. All statistical testing was performed in MATLAB (The MathWorks, Inc., Natick, MA, United States).

#### 1.2.6.2 Post Hoc Analysis

While all younger adults were naïve to split-belt walking, all older adults were experienced at split-belt walking (they had participated in 2-4 split-belt experiments prior to this study), which we did not expect to influence the motor outcome measures of older adults based on our previous work (Sombric et al., 2017). However, we found that several aspects of the motor performance in older, experienced individuals were better than in the younger, naïve participants. Thus, we considered the possibility that exposure to the split-belt condition affected the motor performance of older subjects. To test this idea, we compared the motor outcome measures of older adults when they were naïve versus when they had experienced the split-belt task. Specifically, we used a Wilcoxon signed rank test (paired analysis) to compare the motor outcome measures of Naïve, Older Adults (Old_naïve_) vs. Experienced, Older Adults (Old_experienced_). Of note, only eight (n=8, #women=4, 75.0±2.4 years old) of the eleven participants were included in this analysis because their initial split-belt experiences matched the protocol used in this study (i.e., all protocols had similar belt speed differences introduced abruptly). Cohen’s d effect sizes (e.s.) were computed for each of these statistics. Subjects’ cognitive tasks were not performed during the initial exposures to the split-belt protocols. Thus, future studies are needed to determine the potential effect of exposure on cognitive outcome measures.

## 1.3 Results

### 1.3.1 Old_experienced_ adapt as fast as Young_naïve_ during split-belt walking

We found that older adults can adapt just as fast as younger adults. Figure 2A shows the evolution of locomotor parameters during the Adaptation epoch relative to Baseline behavior. Qualitatively, experienced, older adults (Old_experienced_) adapted Step Length Asymmetry (StepAsym) faster compared to when they were naïve (Old_naïve_), but slower than the naïve, younger adults (Young_naïve_). These differences were quantified with the average Rate of Adaptation (τ) illustrated in the top panel of Figure 2B, where it can be seen that there is a trending effect of exposure (p=0.07, e.s.=0.79) and age (p=0.076, e.s.=0.78). These exposure-related differences in τ for StepAsym were evident in the adaptation of the spatial control of the limb. Specifically, Old_naïve_ adapted their StepPosition (Figure 2A, middle panel) slower than when they were experienced and slower than Young_naïve_, StepTime (Figure 2A, bottom panel). Consistently, the average τ of StepPosition was affected by exposure (Figure 2B, p=0.008, e.s.=0.86), such that Old_experienced_ adapted faster than when they were naïve; and at the same rate as Young_naïve_ (p=0.29, e.s.=0.38). On the other hand, we did not find a significant effect of exposure (p=0.16, e.s.=0.88) or age (p=0.094, e.s.=0.97) on the average τ of StepTime. Note that most of the individual τ values for every subject are identified before the first resting break (i.e., τ <150), therefore the temporal stability of subjects’ memories during resting breaks does not influence the τ for most of the participants. In summary, older adults can adjust spatial gait features as quickly as younger adults with practice.

**Figure 2:**
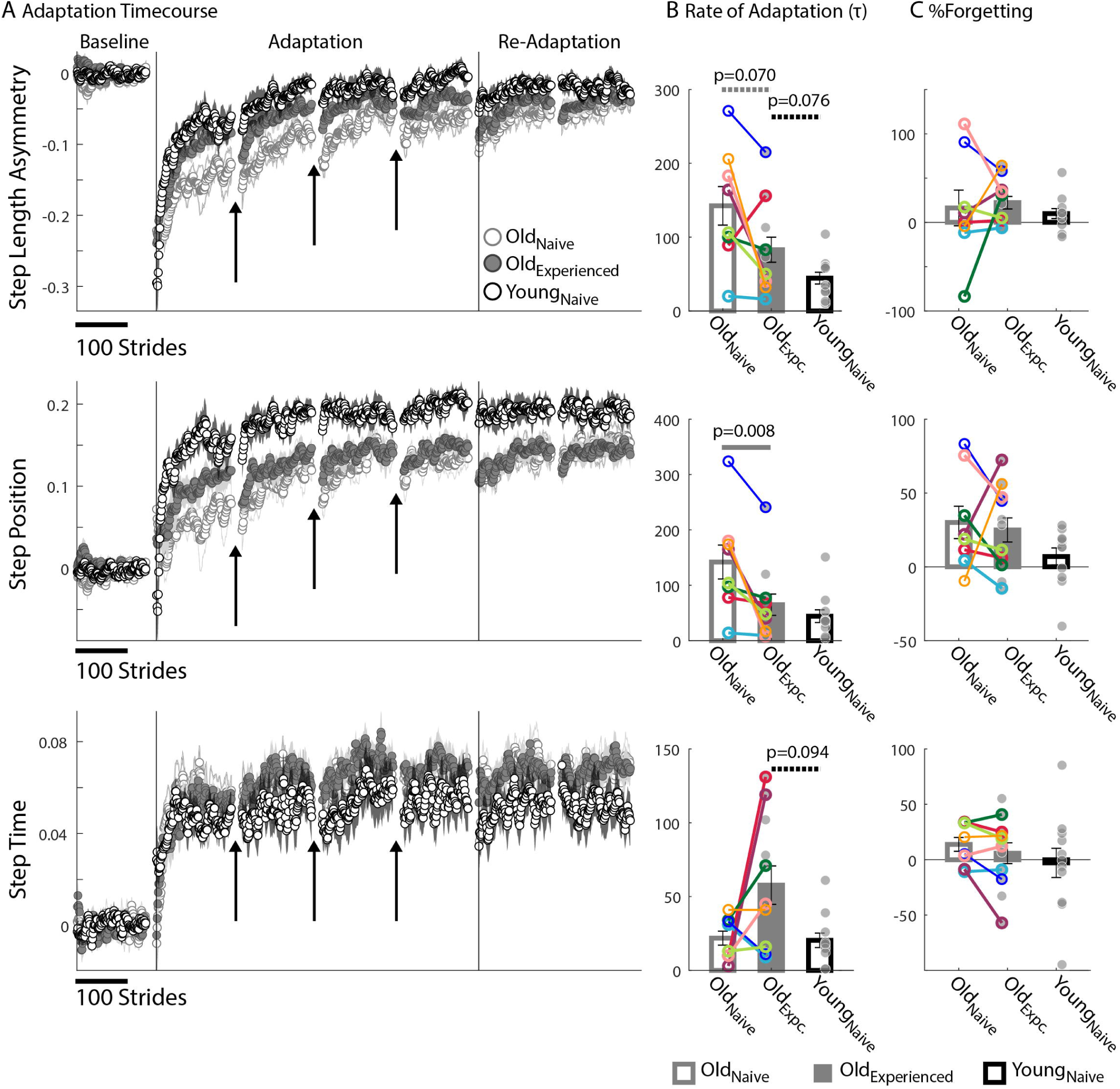
Motor adaptation. (A) Stride-by-stride timecourses of Baseline and Adaptation for StepAsym, StepPosition, and StepTime are illustrated. Black arrows indicate the resting breaks used to compute %Forgetting. Note the decay in the subjects’ adapted state that occurs due to the passage of time during the resting breaks. Dots represent the average of 5 consecutive strides and colored shaded regions indicate the standard error for each group. (B-C) Bar plots indicate the group’s average value ± standard errors. Individual subject behavior is indicated with dots. Dots edged and connected with colored lines (n=8) indicate paired data from the Old_naïve_ and Old_experienced_ testing sessions that were used for statistical testing to determine the effect of exposure. Grey dots (n=11) indicate unpaired data from the Old_experienced_ and Young_naïve_ groups that were used for statistical testing to determine the differences (or lack thereof) between experienced, older adults and naïve, younger adults. (B) Rate of Adaptation (τ): Recall that a larger τ means slower adaptation. The spatial Rate of Adaptation was reduced with exposure. In other words, older adults adapted faster when they had previously experienced the split-belt task compared to when they were naïve. The Rate of Adaptation was not different between the Old_experienced_ and Young_naïve_ adults. (C) %Forgetting: Recall that large positive values of %Forgetting indicated that subjects’ motor states decayed during the resting breaks. Note that there are no age-related differences between groups when the duration of resting breaks was the same between younger and older groups.

### 1.3.2 Motor memories decay with the passage of time similarly across age groups

The split-belt locomotor pattern decayed equally during the resting breaks in younger and older adults. Note in Figure 2A that there are discontinuities in the evolution of the motor adaptation trajectories of all parameters before and after the breaks (shown with arrows) for both older and younger adults. These discontinuities are quantified with %Forgetting illustrated in Figure 2C. Accordingly, we did not find between-group differences in %Forgetting of any gait parameter due to an effect of exposure (StepAsym: p_exposure_=0.74, e.s.=0.26; StepPosition: p_exposure_=0.74, e.s.=0.07; StepTime: p_exposure_=0.31, e.s.=0.37) or age (StepAsym: p_age_=0.26, e.s.=0.54; StepPosition: p_age_=0.21, e.s.=0.70; StepTime: p_age_=0.56, e.s.=0.22). Importantly, we ensured that the resting breaks lasted the same duration in all individuals. Thus, older and younger adults forget similarly motor memories acquired on the split-belt treadmill.

### 1.3.3 Older and younger adults differently adapt spatial and temporal gait features

Older adults counteracted the split-belt perturbation by preferentially adapting temporal, rather than spatial, gait features compared to younger adults. Figure 2A indicates that all groups reached similar Steady States in StepAsym by the end of the Re-Adaptation epoch, but different Steady States in StepPosition (spatial parameter) and StepTime (temporal parameter). These Steady State differences between groups are shown in Figure 3A. There were no differences across groups in the final StepAsym values (p_exposure_=0.95, e.s.=0.12, p_age_=0.43, e.s.=0.36). However, each age group reached similar Steady State StepAsym values with different degrees of spatial and temporal adaptation. Specifically, older adults reached a lower Steady State in StepPosition than younger adults (p_age_=0.030, e.s.=0.28; Figure 3A middle panel); and these Steady State values were comparable in Old_naïve_ and Old_experienced_ testing sessions (p_exposure_=0.64, e.s.=0.97). Conversely, older adults reached higher temporal Steady State values than younger adults (p_age_=0.036, e.s.=0.76; Figure 3A bottom panel); and these Steady State values were similar when older subjects were experienced versus when they were naive (p_exposure_=0.95, e.s.=0.14). In summary, younger and older adults acquired a different stepping pattern on the split-belt treadmill: the younger group exhibited more adaptation of StepPosition and less adaptation of StepTime compared to older adults to reach similar Step Length Asymmetries (StepAsym) during split-belt walking.

**Figure 3.**
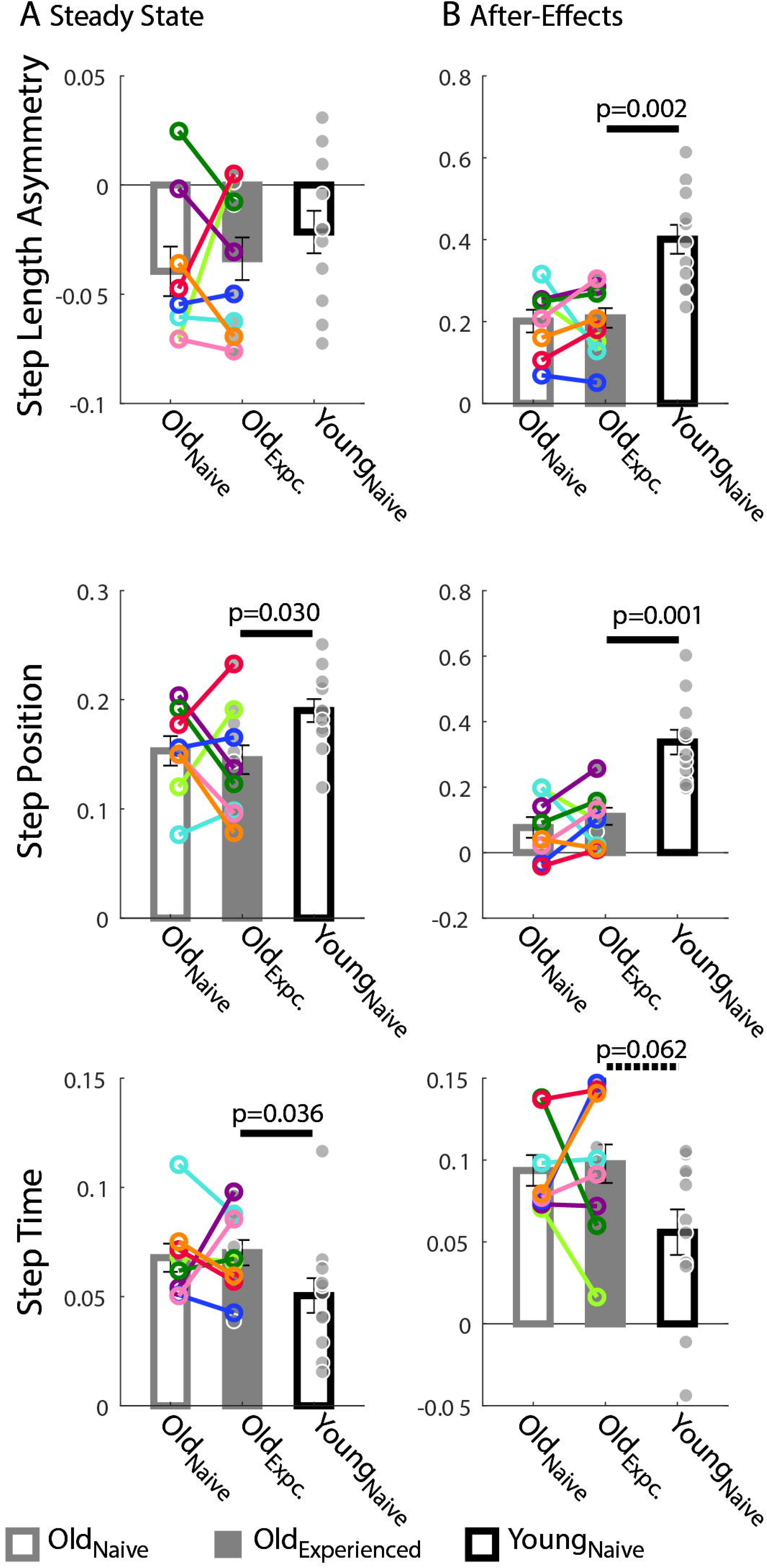
Late Adaptation Behavior and Learning. (A-B) Bar plots indicate the group mean value ± standard errors. Individual subject behavior is indicated with dots. Dots edged and connected with colored lines (n=8) indicate paired data from the Old_naïve_ and Old_experienced_ testing sessions that were used for statistical testing to determine the effect of exposure. Grey dots (n=11) indicate unpaired data from the Old_experienced_ and Young_naïve_ groups that were used for statistical testing to determine if older adults could behave like naïve, younger adults with exposure. (A) Steady State: While all groups reached similar Step Length Asymmetry by the end of the Adaptation period, the spatial and temporal acquired patterns were different between older and younger subjects. Older adults adapted their StepTime more, whereas younger adults adapted their StepPosition more. (B) After-Effects: Large After-Effect values indicate greater change in the gait pattern acquired on the split-belt treadmill. Consistent with the Adaptation phase, older adults had greater StepTime adaptation effects, whereas younger participants exhibited more StepPosition adaptation.

Consistent with the Steady State results, older adults had smaller spatial and larger temporal After-Effects on the treadmill. Treadmill After-Effects quantified during the Catch epoch are illustrated in Figure 3C. StepAsym and StepPosition After-Effects were both significantly smaller for Old_experienced_ than Young_naïve_ (StepAsym: p_age_=0.002, e.s.=1.34; StepPosition, p_age_=0.001, e.s.=1.41). On the other hand, StepTime After-Effects were somewhat larger for Old_experienced_ than Young_naïve_ (p_age_=0.062, e.s.=0.86). Lastly, After-Effects were not different in any of the parameters in Old_experienced_ compared to when they were naïve (Old_naïve_). This is shown by the non-significant exposure effect on After-Effects in all parameters (StepAsym: p_exposure_=0.74, e.s.=0.046; StepPosition: p_exposure_=0.55, e.s.=0.25; StepTime: p_exposure_=0.64, e.s.=0.07). In summary, the same pattern of between-group differences is observed during (Steady State) and after (After-Effects) split-belt walking: younger adults have larger Treadmill After-Effects in StepPosition and smaller After-Effects in StepTime compared to older adults.

### 1.3.4 Spatial and temporal motor perseveration are differently influenced by age and exposure

We found that older adults have more difficulty switching between temporal than spatial motor patterns compared to younger participants. This is shown by the relatively larger Motor Perseveration Errors in older adults compared to young subjects when walking overground in StepTime compared to StepPosition. Figure 4A shows the evolution of StepPosition (spatial parameter) and StepTime (temporal parameter) during overground walking. Note that following Adaptation, participants exhibited asymmetries in StepTime and StepPosition that were different than those during Baseline overground walking, indicating that individuals had difficulties disengaging the spatial and temporal motor patterns that they acquired on the split-belt treadmill when walking overground. This difficulty to switch back to baseline walking patterns was quantified with two perseveration measures: Motor Perseveration Errors (Figure 4B) and %Motor Perseveration (i.e., Motor Perseveration Errors expressed as a percent of Steady State values; Figure 4C). The %Motor Perseveration measure was computed to consider the group differences during the Adaptation period.

**Figure 4.**
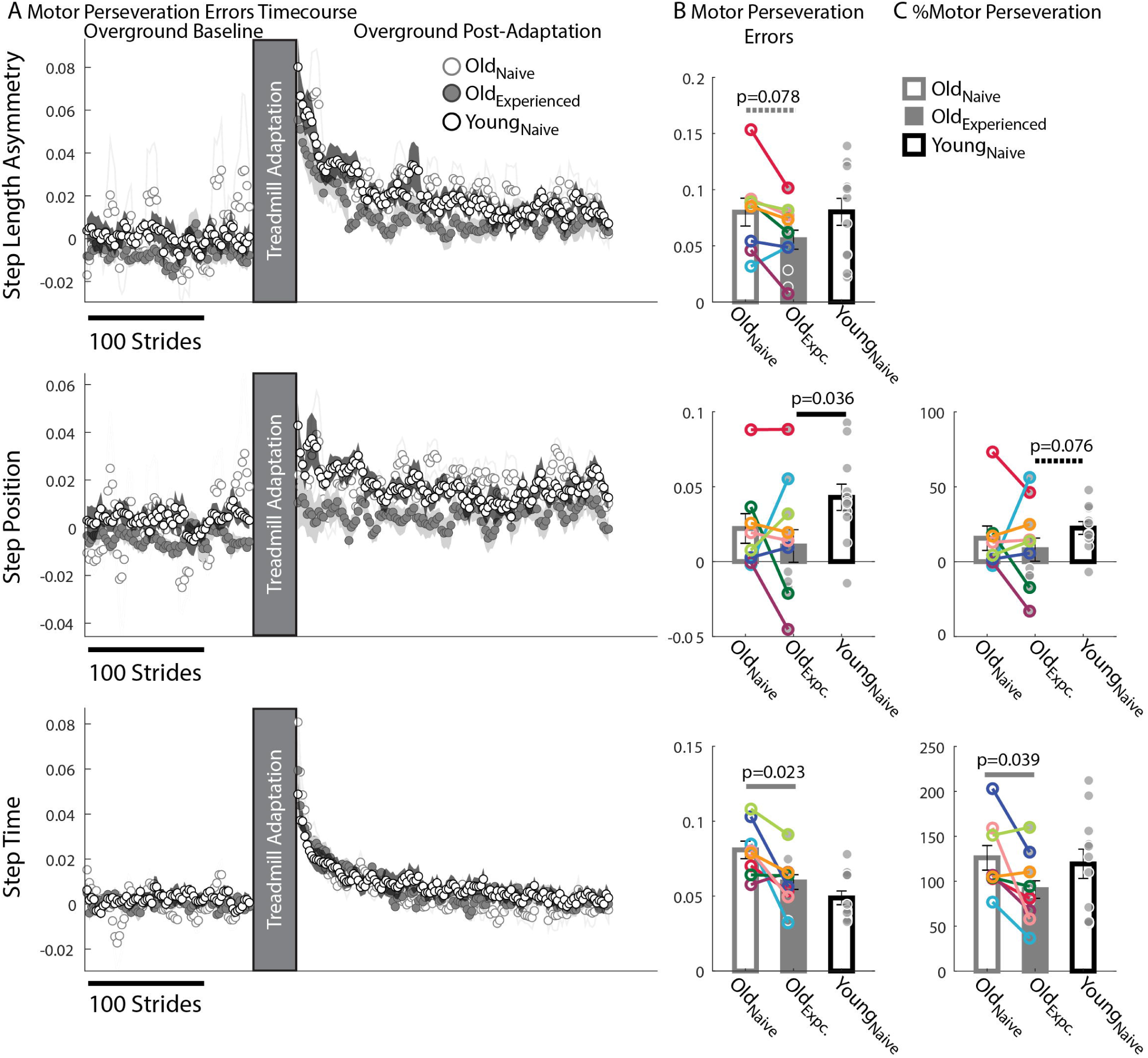
Motor Perseveration During Overground Walking Following Split-Belt Walking. (A) Stride-by-stride timecourses of asymmetries in StepAsym, StepPosition, and StepTime when walking overground during Baseline and Post-Adaptation. Dots represent the average asymmetry values of 5 consecutive strides and colored shaded regions indicate the standard error for each group. (B-C) Bar plots indicate the group’s mean ± standard errors. Individual subject values are indicated with dots. Dots edged and connected with colored lines (n=8) indicate paired data from the Old_naïve_ and Old_experienced_ testing sessions that were used for statistical testing to determine the effect of exposure. Grey dots (n=11) indicate unpaired data from the Old_experienced_ and Young_naïve_ groups that were used for statistical testing to determine if older adults could behave like naïve, younger adults with exposure. (B) Motor Perseveration Errors: Non-zero values indicate that participants cannot disengage the motor pattern acquired on the split-belt treadmill when walking overground. Older adults exhibited greater Motor Perseveration Errors in StepTime than in StepPosition compared to younger adults. (C) %Motor Perseveration: this metric expresses Motor Perseveration Errors as a percent of Steady States to account for the fact that each age group reached different Steady States during split-belt walking. This metric was not computed for StepAsym because Steady States approached zero values.

Motor Perseveration Errors in StepAsym were similar across groups. Notably, we did not observe a significant age effect (p_age_=0.19, e.s.=0.66) in the mean Motor Perseveration Errors of StepAsym. Moreover, we found marginal reductions in Motor Perseveration Errors when older adults were naïve vs. when they were experienced (p_exposure_=0.078, e.s.=0.52). For StepPosition (spatial parameter), we found similar perseveration between older adults when they are naïve vs. experienced (Motor Perseveration Errors: p_exposure_=0.95, e.s.=0.09; %Motor Perseveration p_exposure_=1.00, e.s.=0.07). On the other hand, younger adults exhibited greater Motor Perseveration Errors than older, experienced individuals (p_age_=0.036, e.s.=0.87, Figure 4A middle panel). Of note, this age effect was not observed in the %Motor Perseveration (Figure 4C; %Motor Perseveration: p_age_=0.076, e.s.=0.64), which considers the different Steady States reached during split-belt walking. Thus, age-mediated differences in motor perseveration of StepPosition patterns might be mainly due to differences in the StepPosition pattern acquired on the split-belt treadmill.

The opposite effects were observed in the motor perseveration of timing patterns: we found an exposure effect, and not an age effect, on the perseveration of StepTime. More specifically, motor perseveration of StepTime patterns were smaller in Old_experienced_ than Old_naïve_ groups, as indicated by the significant effect of exposure on both Motor Perseveration Errors (p_expoure_=0.023, e.s.=1.06) and %Motor Perseveration of StepTime (p_exposure_=0.039, e.s.=0.60). Conversely, we did not find age differences in either Motor Perseveration Errors (p_age_=0.24, e.s.=0.62) or %Motor Perseveration (p_age_=0.36, e.s.=0.60) of StepTime. In summary, motor perseveration of spatial patterns was affected by age, whereas motor perseveration of temporal patterns was affected by experience.

### 1.3.5 Cognitive Perseveration is associated with motor perseveration of spatial gait patterns

We found age-related decline in spatial memory and cognitive switching (i.e., large cognitive perseveration errors), but only the latter was associated to motor perseveration in older individuals. Figure 5 shows the larger cognitive perseveration errors in the Old_experienced_ group compared to younger participants in the Young_naïve_ group (Figure 5A p=0.054, e.s.=0.81) in the modified Wisconsin Card Sorting Task. We also found poorer accuracy in the spatial working memory task of older adults compared to younger adults (p=0.032, e.s.=0.94; Figure 5C). Note that we could only perform these analyses between Old_experienced_ and Young_naïve_ groups because cognitive measures where not collected in older individuals when they were naïve to the motor task (Old_naïve_ group). Interestingly, the older subjects’ difficulty to switch actions in the cognitive task were associated to motor switching in the locomotor task. More specifically, older adults with larger Cognitive Perseveration Errors (i.e., poorer cognitive switching), also exhibited larger spatial Motor Perseveration Errors in the locomotor task (Spearman’s correlation: rho=0.71, p=0.014, significant linear correlation: p=0.003, R^2=0.64; 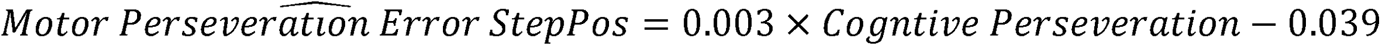. This association between cognitive and motor perseveration was not observed for StepTime (Spearman’s correlation: p=0.74), which is a gait parameter more implicitly controlled (Malone and Bastian, 2010) and it was also exclusive to older individuals (StepPosition Spearman’s correlation: p=0.25; StepTime Spearman’s correlation: p=0.49 in young adults). Lastly, the relation between performance in the cognitive and motor domains is exclusive to cognitive tasks that assess switching ability, rather than age-related cognitive decline in general. Namely, the accuracy of spatial working memory was not associated to motor perseveration errors of any age group (Figure 5D; Older, StepPosition: Spearman’s correlation: p=0.40; Younger, StepPosition: Spearman’s correlation: p=0.95; Older, StepTime: Spearman’s correlation: p=0.99; Younger, StepTime: Spearman’s correlation: p=0.11). In summary, cognitive and motor perseveration become related with healthy aging for motor aspects that are more explicitly controlled in locomotion, such as “where” to step (StepPosition).

**Figure 5.**
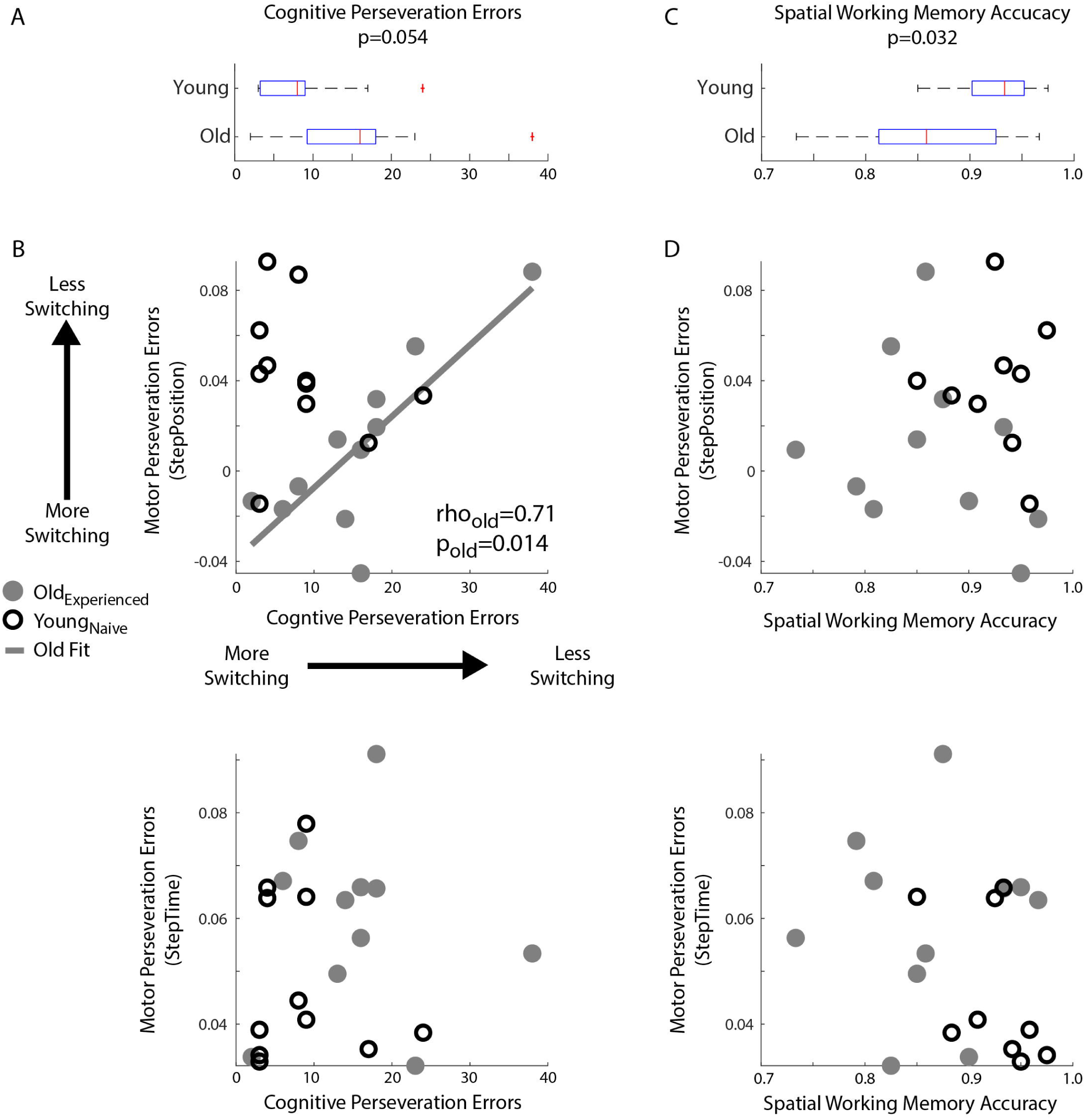
Relationship between Motor and Cognitive Perseveration. (A, B) Box plots indicated age differences in (A) Cognitive Perseveration Errors accrued during the card matching task and (B) Spatial Working Memory Accuracy from the spatial working memory task. The middle, red line indicates the median values for the group, blue edges to the box represent the 25^th^ and 75^th^ percentiles. The whiskers indicate the range of the data excluding outliers, and outliers are shown as red markers. (C, D) Scatter plots indicate the relationship between perseveration errors in motor and cognitive tasks. A linear regression line was displayed when we found significant Spearman’s coefficients (B) Perseveration Errors: larger values indicate greater motor (y-axis) or cognitive (x-axis) perseveration errors. Note a positive association between cognitive and motor perseveration errors in older (grey circles), but not younger adults (empty black circles). (D) Spatial Working Memory Accuracy: There is no relationship between Spatial Working Memory Accuracy and Motor Perseveration Errors for either older or younger subjects.

## 1.4 Discussion

### 1.4.1 Summary

We tested the hypothesis that cognitive-mediated processes for switching are recruited to transition between motor patterns in older adults. To this end, young and older adults adapted their walking pattern on a split-belt treadmill and we measured the motor perseveration of spatial patterns (i.e., StepPosition) and timing patterns (i.e., StepTime) during walking transitions from the split-belt treadmill to overground. We also measured cognitive perseveration with a card matching task in which participants had to switch between different matching rules. We found that cognitive perseveration was associated to motor perseveration in older, but not younger adults. We also found this association was significant in the motor perseveration of the spatial motor pattern, but not the temporal one. These results support the idea that cognitive processes are recruited to compensate for age-related decline in motor switching, but this compensation only benefits motor aspects that are usually controlled explicitly. We also observed that younger and older adults differently adapted spatial and temporal patterns on the split-belt treadmill, but forget them equally during resting breaks. Lastly, a post hoc analysis indicated that many exposures to split-belt walking reduced motor perseveration and increased the rate of adaptation in older individuals compared to when they were naïve to the split-belt task. Taken together, our findings indicate the extent to which cognitive-mediated switching and practice can help older adults regain motor switching abilities similar to those of younger individuals.

### 1.4.2 Cognitive processes compensate for motor perseveration in older adults

We found that cognitive and motor perseveration are related in older individuals, suggesting a compensation strategy for age-related decline in motor switching. However, this compensation mechanism appears to be only beneficial for motor switching between patterns explicitly controlled, such as StepPosition, but not between patterns more implicitly controlled, such as StepTime. The differences we see in motor perseveration between healthy young and older adults are largely consistent with age-related changes in the basal ganglia. Namely, motor switching is regulated by the basal ganglia (Balser et al., 2014; Brown and Almeida, 2011; Leunissen et al., 2013), but older adults are known to recruit cognitive processes (Coxon et al., 2010; Vandevoorde and Orban de Xivry, 2019) to compensate for age-related decline in the basal ganglia’s structure (e.g., striatum volume loss) (Wolpe et al., 2020) and function (Bäckman et al., 2006; Ota et al., 2006; Walhovd et al., 2011). Thus, the degeneration of the basal ganglia with healthy aging may force older adults to recruit cognitive resources to switch between motor patterns. Given that automaticity of gait is reduced with healthy aging (e.g., Guimaraes and Isaacs, 1980), it seems reasonable that cognitive compensation would be effective to improve motor switching. However, the reliance on cognitive switching ability impairs implicit motor switching ability (Boyd and Winstein, 2004; Inzelberg et al., 2001; Sombric et al., 2017), which suggests that cognitive compensation strategies may not always be beneficial to motor performance. We find that the adaptation and perseveration of spatial and temporal motor patterns are differently influenced by aging and exposure and they are also differently influenced by cognitive processes as shown by this work and our previous study (Sombric et al., 2017), adding further evidence of the distinct control of movement in the spatial and temporal domains of locomotion.

### 1.4.3 Older adults more easily update temporal, rather than spatial, gait patterns

We observed age-specific differences in the adaptation of spatial and temporal gait patterns. Consistent with our previous work (Sombric et al., 2017), we observed that older and younger adults can similarly counteract the split-belt perturbation as indicated by Step Length Asymmetry Steady State. However, unlike our previous work, we observe age-related differences in the extent to which they adapt spatial versus temporal patterns compared to young adults. These distinct patterns of adaptation between spatial and temporal gait features during split-belt walking are also indicated by the distinct After-Effects in these two domains. Other studies have also reported age-related differences in the Steady State pattern adopted by older adults on the split-belt treadmill (Bruijn et al., 2012; Vandevoorde and Orban de Xivry, 2019; Vervoort et al., 2019). We had possibly not seen this before (Sombric et al., 2017) due to methodological differences. Namely, we used non-parametric statistical analysis, unlike the previous studies (Sombric et al., 2017), because our sample groups were not normally distributed. Also, we used a different approach to compute subject bias, which could have influenced our results. One interpretation for why older adults exhibit more adaptation of timing parameters, rather than spatial ones compared to younger individuals is that older individuals prioritize stability (e.g., Mian et al., 2006), which is more related to timing features (Finley et al., 2013), over efficiency, which is more related to spatial features (Sánchez et al., 2019). Alternatively, older adults might have reduced adaptation of spatial gait features compared to young individuals because these are more explicitly controlled (Matthis et al., 2017) and the adaptation of cognitive constructs of motor tasks decline with healthy aging (Vandevoorde and Orban de Xivry, 2019; Wolpe et al., 2020).

### 1.4.4 Motor perseveration in older adults is reduced with practice transitioning between motor patterns

We observe that motor perseveration in older individuals is reduced with practice. We have previously found that naïve, older individuals exhibited greater motor perseveration compared to younger counterparts when transitioning across walking environments (Sombric et al., 2017). This is consistent with other studies showing larger motor perseveration when switching between trained and untrained situations (e.g., Bock and Girgenrath, 2006; Fernández-Ruiz et al., 2000; Heuer and Hegele, 2008). However, our current results indicate that motor perseveration is reduced in older adults who have experienced the split-belt protocol before. Therefore, practice transitioning between motor patterns could potentially reduce motor perseveration in older populations. Alternatively, we noticed that older adults exhibited less motor perseveration in motor patterns that were less adapted. Thus, it might also be possible that After-Effects were smaller when transitioning from the treadmill to overground because older individuals have greater resistance to adopting the novel split-belt situation as the “new normal” (Iturralde and Torres-Oviedo, 2019).

In other words, another sign of motor perseveration in older adults might be the resistance to updating movements upon new motor demands. Previous work has shown that older adults are resistant to updating their movements as indicated by the reduced Rates of Adaptation (e.g., Anguera et al., 2012; Bruijn et al., 2012; Rodrigue et al., 2005; Trewartha et al., 2014), lack of savings (Bierbaum et al., 2011), and lower Steady States (e.g., Bruijn et al., 2012; Hegele and Heuer, 2013, 2010; Huang and Ahmed, 2014; Langan and Seidler, 2011; McNay and Willingham, 1998; Seidler, 2006; Wolpe et al., 2020). We considered that forgetting of motor memories could contribute to older adults’ resistance to updating movements (Krishnan et al., 2018; Malone and Bastian, 2016; Sombric et al., 2017). However, we believe this is not the case since younger and older subjects exhibit the same extent of forgetting when the duration of resting breaks is the same between age groups. Thus we consider that older adults’ resistance to updating movement (i.e., motor perseveration) could indicate a higher reliance on previous experiences (Wolpert et al., 1995) due to the larger age-related uncertainty of sensory signals (Goble et al., 2009; Maheu et al., 2015; Rand et al., 2013; Zhang et al., 2008) and motor noise (Holloszy and Larsson, 1995; Kallio et al., 2012; Laidlaw et al., 2000; Vanden Noven et al., 2014; Wolpe et al., 2016).

Nonetheless repeated exposure to the motor tasks makes older adults less resistant to updating movements. This is indicated not only by the reduced motor perseveration when transitioning between walking environments but also by faster adaptation rates in experienced, older adults compared to naïve, older adults. This is consistent with previous work showing that practice improves the Rate of Adaptation (Bock and Schneider, 2002; Pavol et al., 2004) and reduced motor perseveration (Walter et al., 2019). As a side note, previous exposures did not improve forgetting (Sombric et al., 2017), indicating that this might be mediated by different neural processes than motor perseveration. Alternatively, the lack of exposure effects on %Forgetting could be due to the competing mechanisms of aging and practice. In other words, aging could increase forgetting in older individuals, but repeated exposure to the split-belt task could have reduced this forgetting effect. Future studies should test the effect of exposure within a shorter time scale than our study to disentangle the effect of age-related decline and exposure.

### 1.4.5. Clinical Implications

Switching between motor patterns is of clinical interest because of its potential impact on fall prevention. Older adults are at a higher risk of falling possibly due to their difficulty transitioning between walking patterns shaped for distinct environmental demands (Bunterngchit, Yuthachai, Thurmon Lockhart, Jeffrey C. Woldstad, 2000; Lockhart et al., 2002; Lockhart, 1997). Our findings are promising because they indicate that older adults can improve age-related deficits in switching between motor patterns with practice despite reduced neural resources. Consistently, previous studies have demonstrated that exposure to different walking environments reduces the probability of falls in older adults (Hornbrook et al., 1994; Tinetti, 1994; Tinetti et al., 1996; Wagner et al., 1994). Our results, taken together with our previous study (Sombric et al., 2017), indicate that cognitive-mediated processes for switching are recruited to transition between motor patterns in older individuals, but that this is only beneficial when said patterns are controlled explicitly. Thus, interventions that train individuals to use cognitive resources to switch motor patterns may effectively improve spatial aspects of locomotion. Of note, an important limitation of our study is that the relation between perseveration in the cognitive and motor domains was identified in older adults who had previously experienced the split-belt task multiple times. Therefore, future studies are needed to determine if cognitive strategies are developed with practice switching between motor patterns. In sum, our work highlights the importance of exposure and cognitive compensation to reduce motor perseveration in older individuals, which could be used to decrease fall risk in the elderly.

## 1.5 Conclusion

In conclusion, we found that cognitive resources and practice can regulate motor perseveration in older individuals. Our results are novel because we observed an interesting correlation between cognitive and motor perseveration in older individuals, suggesting that processes mediating switching actions might be unified in these two domains as we age. Moreover, we found that older subjects who have practiced switching between different walking patterns can achieve similar motor perseveration and forgetting of context-specific motor memories to those observed in naïve, younger individuals. Importantly, older and younger subjects adapt spatial and temporal aspects of gait differently, which could influence the degree of After-Effects (or motor perseveration) that is observed on and off the treadmill. Taken together, our results are important because they indicate that deficient motor switching in older populations can be improved through practice and cognitive compensation.

## Availability of Data and Materials

The datasets analyzed during the current study and corresponding code are available at https://osf.io/yrca5/.

## Acknowledgments

The authors thank Dr. Andrea Weinstein for her insightful feedback on the data analysis of the cognitive tasks. The authors thank Megan Thorbahn for her dedication to data collection and data processing.

## References

Albergaria, C., Silva, N.T., Pritchett, D.L., Carey, M.R., Carey, M.R., 2018. Locomotor activity modulates associative learning in mouse cerebellum. Nat. Neurosci. 21. https://doi.org/10.1038/s41593-018-0129-x

Anguera, J. a., Bernard, J. a., Jaeggi, S.M., Buschkuehl, M., Benson, B.L., Jennett, S., Humfleet, J., Reuter-Lorenz, P. a., Jonides, J., Seidler, R.D., 2012. The effects of working memory resource depletion and training on sensorimotor adaptation. Behav. Brain Res. 228, 107–115. https://doi.org/10.1016/j.bbr.2011.11.040

Anguera, J. a, Reuter-Lorenz, P. a, Willingham, D.T., Seidler, R.D., 2011. Failure to engage spatial working memory contributes to age-related declines in visuomotor learning. J. Cogn. Neurosci. 23, 11–25. https://doi.org/10.1162/jocn.2010.21451

Bäckman, L., Nyberg, L., Lindenberger, U., Li, S.-C., Farde, L., 2006. The correlative triad among aging, dopamine, and cognition: Current status and future prospects. Neurosci. Biobehav. Rev. 30, 791–807. https://doi.org/10.1016/j.neubiorev.2006.06.005

Balser, N., Lorey, B., Pilgramm, S., Stark, R., Bischoff, M., Zentgraf, K., Williams, A.M., Munzert, J., 2014. Prediction of human actions: Expertise and task-related effects on neural activation of the action observation network. Hum. Brain Mapp. 35, 4016–4034. https://doi.org/10.1002/hbm.22455

Bierbaum, S., Peper, A., Karamanidis, K., Arampatzis, A., 2011. Adaptive feedback potential in dynamic stability during disturbed walking in the elderly. J. Biomech. 44, 1921–1926. https://doi.org/10.1016/j.jbiomech.2011.04.027

Bo, J., Borza, V., Seidler, R.D., 2009. Age-related declines in visuospatial working memory correlate with deficits in explicit motor sequence learning. J. Neurophysiol. 102, 2744–2754. https://doi.org/10.1152/jn.00393.2009

Bock, O., Girgenrath, M., 2006. Relationship between sensorimotor adaptation and cognitive functions in younger and older subjects. Exp. brain Res. 169, 400–406. https://doi.org/10.1007/s00221-005-0153-4

Bock, O., Schneider, S., 2002. Sensorimotor adaptation in young and elderly humans. Neurosci. Biobehav. Rev. 26, 761–767. https://doi.org/10.1016/S0149-7634(02)00063-5

Boone, K.B., Ghaffarian, S., Lesser, I.M., Hill-Gutierrez, E., Berman., N.G., 1993. Wisconsin card sorting test performance in healthy, older adults: relationship to age, sex, education, and IQ. J. Clin. Psychol. 49, 54–60.

Boyd, L. a., Winstein, C.J., 2004. Providing Explicit Information Disrupts Implicit Motor Learning After Basal Ganglia Stroke. Learn. Mem. 11, 388–396. https://doi.org/10.1101/lm.80104

Brown, M.J.N., Almeida, Q.J., 2011. Evaluating dopaminergic system contributions to cued pattern switching during bimanual coordination. Eur. J. Neurosci. 34, 632–640. https://doi.org/10.1111/j.1460-9568.2011.07773.x

Bruijn, S.M., Van Impe, A., Duysens, J., Swinnen, S.P., 2012. Split-belt walking: adaptation differences between young and older adults. J. Neurophysiol. 108, 1149–57. https://doi.org/10.1152/jn.00018.2012

Bunterngchit, Yuthachai, Thurmon Lockhart, Jeffrey C. Woldstad, and J.L.S., 2000. Age related effects of transitional floor surfaces and obstruction of view on gait characteristics related to slips and falls. Int. J. Ind. Ergon. 25, 223–232. https://doi.org/10.1158/0008-5472.CAN-10-4002.BONE

Christou, A.I., Miall, R.C., McNab, F., Galea, J.M., 2016. Individual differences in explicit and implicit visuomotor learning and working memory capacity. Sci. Rep. 6, 1–13. https://doi.org/10.1038/srep36633

Coubard, O.A., Duretz, S., Lefebvre, V., Lapalus, P., Ferrufino, L., Heath, J.E., 2011. Practice of contemporary dance improves cognitive flexibility in aging. Front. Aging Neurosci. 3, 13. https://doi.org/10.3389/fnagi.2011.00013

Coxon, J.P., Goble, D.J., Van Impe, A., De Vos, J., Wenderoth, N., Swinnen, S.P., 2010. Reduced basal ganglia function when elderly switch between coordinated movement patterns. Cereb. Cortex 20, 2368–2379. https://doi.org/10.1093/cercor/bhp306

Daigneault, S., Braun, C.M., Whitaker., H.A., 1992. Early effects of normal aging on perseverative and non-perseverative prefrontal measures. Dev. Neuropsychol. 8, 99–114.

Eppinger, B., Hämmerer, D., Li, S.-C., 2011. Neuromodulation of reward-based learning and decision making in human aging. Ann. N. Y. Acad. Sci. 1235, 1–17. https://doi.org/10.1111/j.1749-6632.2011.06230.x.Neuromodulation

Erickson, K.I., Prakash, R.S., Voss, M.W., Chaddock, L., Hu, L., Morris, K.S., White, S.M., Wójcicki, T.R., McAuley, E., Kramer, A.F., 2009. Aerobic fitness is associated with hippocampal volume in elderly humans. Hippocampus 19, 1030–1039. https://doi.org/10.1002/hipo.20547

Erickson, K.I., Voss, M.W., Prakash, R.S., Basak, C., Szabo, A., Chaddock, L., Kim, J.S., Heo, S., Alves, H., White, S.M., Wojcicki, T.R., Mailey, E., Vieira, V.J., Martin, S. a, Pence, B.D., Woods, J. a, McAuley, E., Kramer, A.F., 2011. Exercise training increases size of hippocampus and improves memory. Proc. Natl. Acad. Sci. U. S. A. 108, 3017–3022. https://doi.org/10.1073/pnas.1015950108

Erickson, K.I., Weinstein, A.M., Sutton, B.P., Prakash, R.S., Voss, M.W., Chaddock, L., Szabo, A.N., Mailey, E.L., White, S.M., Wojcicki, T.R., McAuley, E., Kramer, A.F., 2012. Beyond vascularization: Aerobic fitness is associated with N-acetylaspartate and working memory. Brain Behav. 2, 32–41. https://doi.org/10.1002/brb3.30

Fernández-Ruiz, J., Hall, C., Vergara, P., Díaz, R., 2000. Prism adaptation in normal aging: Slower adaptation rate and larger aftereffect. Cogn. Brain Res. 9, 223–226. https://doi.org/10.1016/S0926-6410(99)00057-9

Finley, J.M., Bastian, A.J., Gottschall, J.S., 2013. Learning to be economical: the energy cost of walking tracks motor adaptation. J. Physiol. 591, 1081–1095. https://doi.org/10.1113/jphysiol.2012.245506

Finley, J.M., Long, A., Bastian, A.J., Torres-Oviedo, G., 2015. Spatial and Temporal Control Contribute to Step Length Asymmetry During Split-Belt Adaptation and Hemiparetic Gait. Neurorehabil Neural Repair 29, 786–795. https://doi.org/1545968314567149

Goble, D.J., Coxon, J.P., Wenderoth, N., Van Impe, A., Swinnen, S.P., 2009. Proprioceptive sensibility in the elderly: Degeneration, functional consequences and plastic-adaptive processes. Neurosci. Biobehav. Rev. 33, 271–278. https://doi.org/10.1016/j.neubiorev.2008.08.012

Guimaraes, R.M., Isaacs, B., 1980. Characteristics of the gait in old people who fall. Int. Rehabil. Med. 2, 177–180.

Haaland, Y.K., Vranes, L.F., Goodwin, J.S., Garry, P.J., 1987. Wisconsin Card Sort Test performance in a healthy elderly population. J. Gerontol. 42, 345–346.

Head, D., Kennedy, K.M., Rodrigue, K.M., Raz, N., 2009. Age differences in perseveration: Cognitive and neuroanatomical mediators of performance on the Wisconsin Card Sorting Test. Neuropsychologia 47, 1200–1203. https://doi.org/10.1016/j.neuropsychologia.2009.01.003

Hegele, M., Heuer, H., 2013. Age-related variations of visuomotor adaptation result from both the acquisition and the application of explicit knowledge. Psychol. Aging 28, 333–9. https://doi.org/10.1037/a0031914

Hegele, M., Heuer, H., 2010. Adaptation to a direction-dependent visuomotor gain in the young and elderly. Psychol. Res. 74, 21–34. https://doi.org/10.1007/s00426-008-0221-z

Heuer, H., Hegele, M., 2008. Adaptation to visuomotor rotations in younger and older adults. Psychol. Aging 23, 190–202. https://doi.org/10.1037/0882-7974.23.1.190

Holloszy, J.O., Larsson, L., 1995. Motor units: remodeling in aged animals. J Gerontol A Biol Sci Med Sci 50, 91–95.

Hornbrook, M.C., Stevens, V.J., Wingfield, D.J., Hollis, J.F., Greenlick, M.R., Ory, M.G., 1994. Preventing falls among community-dwelling older persons: results from a randomized trial. Gerontologist 34, 16–23.

Hötting, K., Holzschneider, K., Stenzel, A., Wolbers, T., Röder, B., 2013. Effects of a cognitive training on spatial learning and associated functional brain activations. BMC Neurosci. 14, 73. https://doi.org/10.1186/1471-2202-14-73

Huang, H.J., Ahmed, A.A., 2014. Older adults learn less, but still reduce metabolic cost, during motor adaptation. J. Neurophysiol. 111, 135–44. https://doi.org/10.1152/jn.00401.2013

Inzelberg, R., Plotnik, M., Flash, T., Schechtman, E., Shahar, I., Korczyn, a D., 2001. Mental and motor switching in Parkinson’s disease. J. Mot. Behav. 33, 377–385. https://doi.org/10.1080/00222890109601921

Iturralde, P.A., Torres-Oviedo, G., 2019. Corrective Muscle Activity Reveals Subject-Specific Sensorimotor Recalibration. Eneuro 6, ENEURO.0358-18.2019. https://doi.org/10.1523/eneuro.0358-18.2019

Kallio, J., Søgaard, K., Avela, J., Komi, P., Selänne, H., Linnamo, V., S??gaard, K., Avela, J., Komi, P., Sel??nne, H., Linnamo, V., 2012. Age-related decreases in motor unit discharge rate and force control during isometric plantar flexion. J. Electromyogr. Kinesiol. 22, 983–989. https://doi.org/10.1016/j.jelekin.2012.05.009

Kray, J., Lindenberger, U., 2000. Adult age differences in task switching. Psychol. Aging 15, 126–147. https://doi.org/10.1037//OS82-797-U5.1.126

Krishnan, C., Washabaugh, E.P., Reid, C.E., Althoen, M.M., Ranganathan, R., 2018. Learning new gait patterns: Age-related differences in skill acquisition and interlimb transfer. Exp. Gerontol. 111, 45–52. https://doi.org/10.1016/j.exger.2018.07.001

Laidlaw, D.H., Bilodeau, M., Enoka, R.M., 2000. Steadiness is reduced and motor unit discharge is more variable in old adults. Muscle and Nerve 23, 600–612. https://doi.org/10.1002/(SICI)1097-4598(200004)23:4<600::AID-MUS20>3.0.CO;2-D

Langan, J., Seidler, R.D., 2011. Age differences in spatial working memory contributions to visuomotor adaptation and transfer. Behav. Brain Res. 225, 160–168. https://doi.org/10.1016/j.bbr.2011.07.014

Leunissen, I., Coxon, J.P., Geurts, M., Caeyenberghs, K., Michiels, K., Sunaert, S., Swinnen, S.P., 2013. Disturbed cortico-subcortical interactions during motor task switching in traumatic brain injury. Hum. Brain Mapp. 34, 1254–1271. https://doi.org/10.1002/hbm.21508

Li, Y., Tian, X., Xiong, Z.Y., Liao, J.L., Hao, L., Liu, G.L., Ren, Y.P., Wang, Q., Duan, L.P., Zheng, Z.X., Quan, W.X., Dong, J., 2016. Performance of the modified mini-mental state examination (3MS) in assessing specific cognitive function in patients undergoing peritoneal dialysis. PLoS One 11, 1–12. https://doi.org/10.1371/journal.pone.0166470

Lockhart, T., Woldstad, J., Smith, J., Ramsey, J., 2002. Effects of age related sensory degradation on perception of floor slipperiness and associated slip parameters. Saf. Sci. 40, 689–703. https://doi.org/10.1016/S0925-7535(01)00067-4.Effects

Lockhart, T.E., 1997. The ability of elderly people to traverse slippery walking surfaces., in: Proceedings of the Human Factors and Ergonomics Society Annual Meeting. SAGE Publications, Sage CA: Los Angeles, CA, pp. 125–129.

Maheu, M., Houde, M.-S., Landry, S.P., Champoux, F., 2015. The Effects of Aging on Clinical Vestibular Evaluations. Front. Neurol. 6, 205.

Malone, L. a, Bastian, A.J., 2010. Thinking about walking: effects of conscious correction versus distraction on locomotor adaptation. J. Neurophysiol. 103, 1954–62. https://doi.org/10.1152/jn.00832.2009

Malone, L.A., Bastian, A.J., 2016. Age-related forgetting in locomotor adaptation. Neurobiol. Learn. Mem. 128, 1–6. https://doi.org/10.1016/j.nlm.2015.11.003

Malone, L.A., Bastian, A.J., 2014. Spatial and Temporal Asymmetries in Gait Predict Split-Belt Adaptation Behavior in Stroke. Neurorehabil. Neural Repair 28, 230–240. https://doi.org/10.1177/1545968313505912

Mariscal, D.M., Iturralde, P.A., Torres-Oviedo, G., 2020. Altering attention to split-belt walking increases the generalization of motor memories across walking contexts. J. Neurophysiol. 123, 1838–1848. https://doi.org/10.1152/jn.00509.2019

Matthis, J.S., Barton, S.L., Fajen, B.R., 2017. The critical phase for visual control of human walking over complex terrain. Proc. Natl. Acad. Sci. U. S. A. 114, E6720–E6729. https://doi.org/10.1073/pnas.1611699114

McAuley, E., Szabo, A.N., Mailey, E.L., Erickson, K.I., Voss, M., White, S.M., Wójcicki, T.R., Gothe, N., Olson, E.A., Mullen, S.P., Kramer, A.F., 2011. Non-Exercise Estimated Cardiorespiratory Fitness: Associations with Brain Structure, Cognition, and Memory Complaints in Older Adults. Ment. Health Phys. Act. 4, 5–11. https://doi.org/10.1158/0008-5472.CAN-10-4002.BONE

McNay, E.C., Willingham, D.B., 1998. Deficit in learning of a motor skill requiring strategy, but not of perceptuomotor recalibration, with aging. Learn. Mem. 4, 411–420. https://doi.org/10.1101/lm.4.5.411

Mian, O.S., Thom, J.M., Ardigò, L.P., Narici, M. V., Minetti, A.E., 2006. Metabolic cost, mechanical work, and efficiency during walking in young and older men. Acta Physiol. 186, 127–139. https://doi.org/10.1111/j.1748-1716.2006.01522.x

Nelson, H.E., 1976. A modified card sorting test sensitive to frontal lobe defects. Cortex 12, 313–324.

Ota, M., Yasuno, F., Ito, H., Seki, C., Nozaki, S., Asada, T., Suhara, T., 2006. Age-related decline of dopamine synthesis in the living human brain measured by positron emission tomography with L-[β-11 C] DOPA. Life Sci. 79, 730–736. https://doi.org/10.1016/j.lfs.2006.02.017

Pavol, M.J., Runtz, E.F., Pai, Y.-C., 2004. Young and Older Adults Exhibit Proactive and Reactive Adaptations to Repeated Slip Exposure. Journals Gerontol. Ser. A Biol. Sci. Med. Sci. 59, 494–502. https://doi.org/10.1093/gerona/59.5.m494

Rand, M.K., Wang, L., Müsseler, J., Heuer, H., 2013. Vision and proprioception in action monitoring by young and older adults. Neurobiol. Aging 34, 1864–1872. https://doi.org/10.1016/j.neurobiolaging.2013.01.021

Reisman, D.S., Block, H.J., Bastian, A.J., Darcy, S., Block, H.J., Bastian, A.J., 2005. Interlimb coordination during locomotion: what can be adapted and stored? J. Neurophysiol. 94, 2403–15. https://doi.org/10.1152/jn.00089.2005

Reisman, D.S., McLean, H., Keller, J., Danks, K.A., Bastian, A.J., 2013. Repeated split-belt treadmill training improves poststroke step length asymmetry. Neurorehabil. Neural Repair 27, 460–468. https://doi.org/10.1177/1545968312474118

Ridderinkhof, K.R., Span, M.M., Van Der Molen, M.W., 2002. Perseverative behavior and adaptive control in older adults: Performance monitoring, rule induction, and set shifting. Brain Cogn. 49, 382–401. https://doi.org/10.1006/brcg.2001.1506

Rodrigue, K.M., Kennedy, K.M., Raz, N., 2005. Aging and longitudinal change in perceptual-motor skill acquisition in healthy adults. J. Gerontol. B. Psychol. Sci. Soc. Sci. 60, P174–P181.

Salmi, J., Nyberg, L., Laine, M., 2018. Working memory training mostly engages general-purpose large-scale networks for learning. Neurosci. Biobehav. Rev. 93, 108–122. https://doi.org/10.1016/j.neubiorev.2018.03.019

Sánchez, N., Simha, S.N., Donelan, J.M., Finley, J.M., 2019. Taking advantage of external mechanical work to reduce metabolic cost: the mechanics and energetics of split-belt treadmill walking. J. Physiol. 15, JP277725. https://doi.org/10.1113/JP277725

Seidler, R.D., 2006. Differential effects of age on sequence learning and sensorimotor adaptation. Brain Res. Bull. 70, 337–46. https://doi.org/10.1016/j.brainresbull.2006.06.008

Sombric, C.J., Harker, H.M., Sparto, P.J., Torres-oviedo, G., 2017. Explicit Action Switching Interferes with the Context-Specificity of Motor Memories in Older Adults. Front. Aging Neurosci. 9, 40.

Stuss, D.T., Murphy, K.J., Palumbo, C., Levine, B., Hong, J., Hamer, L., Alexander, M.P., Izukawa, D., 2000. Wisconsin Card Sorting Test performance in patients with focal frontal and posterior brain damage: effects of lesion location and test structure on separable cognitive processes. Neuropsychologia 38, 388–402. https://doi.org/10.1016/s0028-3932(99)00093-7

Szabo, A.N., McAuley, E., Erickson, K.I., Voss, M., Prakash, R.S., Mailey, E.L., Wójcicki, T.R., White, S.M., Gothe, N., Olson, E.A., Kramer, A.F., 2011. Cardiorespiratory Fitness, Hippocampal Volume and Frequency of Forgetting in Older Adults. Neuropsychology 25, 545–553. https://doi.org/10.1037/a0022733.Cardiorespiratory

Tinetti, M.E., 1994. Prevention of falls and fall injuries in elderly persons: a research agenda. Prev. Med. (Baltim). 23, 756–762.

Tinetti, M.E., McAvay, G., Claus, E., 1996. Does multiple risk factor reduction explain the reduction in fall rate in the Yale FICSIT Trial? Frailty and Injuries Cooperative Studies of Intervention Techniques. Am. J. Epidemiol. 144, 389–399. https://doi.org/10.1093/oxfordjournals.aje.a008940

Torres-Oviedo, G., Bastian, A.J., 2012. Natural error patterns enable transfer of motor learning to novel contexts. J. Neurophysiol. 107, 346–56. https://doi.org/10.1152/jn.00570.2011

Torres-Oviedo, G., Bastian, A.J., 2010. Seeing Is BelievingLJ: Effects of Visual Contextual Cues on Learning and Transfer of Locomotor Adaptation. J. Neurosci. 30, 17015–17022. https://doi.org/10.1523/JNEUROSCI.4205-10.2010

Trewartha, K.M., Garcia, A., Wolpert, D.M., Flanagan, J.R., 2014. Fast but fleeting: adaptive motor learning processes associated with aging and cognitive decline. J. Neurosci. 34, 13411–21. https://doi.org/10.1523/JNEUROSCI.1489-14.2014

Uresti-Cabrera, L.A., Diaz, R., Vaca-Palomares, I., Fernandez-Ruiz, J., 2015. The effect of spatial working memory deterioration on strategic visuomotor learning across aging. Behav. Neurol. 2015, 17–19. https://doi.org/10.1155/2015/512617

Vanden Noven, M.L., Pereira, H.M., Yoon, T., Stevens, A.A., Nielson, K.A., Hunter, S.K., 2014. Motor variability during sustained contractions increases with cognitive demand in older adults. Front. Aging Neurosci. 6, 97. https://doi.org/10.3389/fnagi.2014.00097

Vandevoorde, K., Orban de Xivry, J.J., 2019. Internal model recalibration does not deteriorate with age while motor adaptation does. Neurobiol. Aging 80, 138–153. https://doi.org/10.1016/j.neurobiolaging.2019.03.020

Vervoort, D., Rob Den Otter, A., Buurke, T.J.W., Vuillerme, N., Hortobágyi, T., Lamoth, C.J.C., 2019. Effects of aging and task prioritization on split-belt gait adaptation. Front. Aging Neurosci. 11, 1–12. https://doi.org/10.3389/fnagi.2019.00010

Volkow, N.D., Gur, R.C., Wang, G.J., Fowler, J.S., Moberg, P.J., Ding, Y.S., Hitzemann, R., Smith, G., Logan, J., 1998. Association between decline in brain dopamine activity with age and cognitive and motor impairment in healthy individuals. Am. J. Psychiatry 155, 344–349. https://doi.org/10.1176/ajp.155.3.344

Voss, M.W., Prakash, R.S., Erickson, K.I., Basak, C., Chaddock, L., Kim, J.S., Alves, H., Heo, S., Szabo, A.N., White, S.M., Wójcicki, T.R., Mailey, E.L., Gothe, N., Olson, E.A., McAuley, E., Kramer, A.F., 2010. Plasticity of brain networks in a randomized intervention trial of exercise training in older adults. Front. Aging Neurosci. 2, 32. https://doi.org/10.3389/fnagi.2010.00032

Wagner, E.H., LaCroix, A.Z., Grothaus, L., Leveille, S.G., Hecht, J.A., Artz, K., Odle, K., Buchner, D.M., 1994. Preventing disability and falls in older adults: a population-based randomized trial. Am. J. Public Health 84, 1800–1806. https://doi.org/10.2105/ajph.84.11.1800

Walhovd, K.B., Westlye, L.T., Amlien, I., Espeseth, T., Reinvang, I., Raz, N., Agartz, I., Salat, D.H., Greve, D.N., Fischl, B., Dale, A.M., Fjell, A.M., 2011. Consistent neuroanatomical age-related volume differences across multiple samples. Neurobiol. Aging 32, 916–932. https://doi.org/10.1016/j.neurobiolaging.2009.05.013

Walter, C.S., Hengge, C.R., Lindauer, B.E., Schaefer, S.Y., 2019. Declines in motor transfer following upper extremity task-specific training in older adults. Exp. Gerontol. 116, 14–19. https://doi.org/10.1016/j.exger.2018.12.012

Weinstein, A.M., Voss, M.W., Prakash, R.S., Chaddock, L., Szabo, A., White, S.M., Wojcicki, T.R., Mailey, E., McAuley, E., Kramer, A.F., Erickson, K.I., 2012. The association between aerobic fitness and executive function is mediated by prefrontal cortex volume. Brain. Behav. Immun. 26, 811–819. https://doi.org/10.1016/j.bbi.2011.11.008

Wolpe, N., Ingram, J.N., Tsvetanov, K.A., Geerligs, L., Kievit, R.A., Henson, R.N., Wolpert, D.M., Rowe, J.B., Tyler, L.K., Brayne, C., Bullmore, E., Calder, A., Cusack, R., Dalgleish, T., Duncan, J., Matthews, F.E., Marslen-Wilson, W., Shafto, M.A., Campbell, K., Cheung, T., Davis, S., McCarrey, A., Mustafa, A., Price, D., Samu, D., Taylor, J.R., Treder, M., van Belle, J., Williams, N., Bates, L., Emery, T., Erzinclioglu, S., Gadie, A., Gerbase, S., Georgieva, S., Hanley, C., Parkin, B., Troy, D., Auer, T., Correia, M., Gao, L., Green, E., Henriques, R., Allen, J., Amery, G., Amunts, L., Barcroft, A., Castle, A., Dias, C., Dowrick, J., Fair, M., Fisher, H., Goulding, A., Grewal, A., Hale, G., Hilton, A., Johnson, F., Johnston, P., Kavanagh-Williamson, T., Kwasniewska, M., McMinn, A., Norman, K., Penrose, J., Roby, F., Rowland, D., Sargeant, J., Squire, M., Stevens, B., Stoddart, A., Stone, C., Thompson, T., Yazlik, O., Barnes, D., Dixon, M., Hillman, J., Mitchell, J., Villis, L., 2016. Ageing increases reliance on sensorimotor prediction through structural and functional differences in frontostriatal circuits. Nat. Commun. 7, 1–11. https://doi.org/10.1038/ncomms13034

Wolpe, N., Ingram, J.N., Tsvetanov, K.A., Henson, R.N., Wolpert, D.M., Tyler, L.K., Brayne, C., Bullmore, E.T., Calder, A.C., Cusack, R., Dalgleish, T., Duncan, J., Matthews, F.E., Marslen-Wilson, W.D., Shafto, M.A., Campbell, K., Cheung, T., Davis, S., Geerligs, L., Kievit, R., McCarrey, A., Mustafa, A., Price, D., Samu, D., Taylor, J.R., Treder, M., van Belle, J., Williams, N., Bates, L., Emery, T., Erzinçlioglu, S., Gadie, A., Gerbase, S., Georgieva, S., Hanley, C., Parkin, B., Troy, D., Auer, T., Correia, M., Gao, L., Green, E., Henriques, R., Allen, J., Amery, G., Amunts, L., Barcroft, A., Castle, A., Dias, C., Dowrick, J., Fair, M., Fisher, H., Goulding, A., Grewal, A., Hale, G., Hilton, A., Johnson, F., Johnston, P., Kavanagh-Williamson, T., Kwasniewska, M., McMinn, A., Norman, K., Penrose, J., Roby, F., Rowland, D., Sargeant, J., Squire, M., Stevens, B., Stoddart, A., Stone, C., Thompson, T., Yazlik, O., Barnes, D., Dixon, M., Hillman, J., Mitchell, J., Villis, L., Rowe, J.B., 2020. Age-related reduction in motor adaptation: brain structural correlates and the role of explicit memory. Neurobiol. Aging 90, 13–23. https://doi.org/10.1016/j.neurobiolaging.2020.02.016

Wolpert, D.M., Ghahramani, Z., Jordan, M.I., 1995. An internal model for sensorimotor integration. Science (80-.). 269, 1880–1882.

Zhang, C., Hua, T., Li, G., Tang, C., Sun, Q., Zhou, P., 2008. Visual function declines during normal aging. Curr. Sci. 95, 1544–1550.

